# Aβ-aggregation-generated blue autofluorescence illuminates senile plaques, complex blood and vascular pathologies in the Alzheimer’s disease

**DOI:** 10.1101/2023.07.06.548042

**Authors:** Hualin Fu, Jilong Li, Chunlei Zhang, Peng Du, Guo Gao, Qiqi Ge, Xinping Guan, Daxiang Cui

**Affiliations:** Institute of Nano Biomedicine and Engineering, School of Sensing Science and Engineering, School of Electronic Information and Electrical Engineering, Shanghai Jiao Tong University, 800 Dongchuan Road, Shanghai, 200240, China; Institute of Marine Equipment, Shanghai Jiao Tong University, 211 Wenjin Road, Shanghai, 200240, China; National Center for Translational Medicine, Shanghai Jiao Tong University, 800 Dongchuan Road, Shanghai, 200240, China; Department of Colorectal Surgery, Xinhua Hospital, School of Medicine, Shanghai Jiao Tong University, Shanghai, China; Department of Automation, Shanghai Jiao Tong University, Shanghai 200240, China; The Key Laboratory of System Control and Information Processing, Ministry of Education, Shanghai 200240, China

**Keywords:** senile plaque, blue autofluorescence, Aβ, Hemoglobin, microaneurysm, Alzheimer’s disease

## Abstract

Senile plaque blue autofluorescence in the Alzheimer’s disease (AD) was discovered around 40 years ago, however, its impact on AD pathology is not fully examined. We analyzed senile plaques with immunohistochemistry and fluorescence imaging on AD brain pathological sections and also the Aβ aggregation process *in vitro* in test tubes. In DAPI or Hoechst staining experiments, the data showed that the nuclear blue fluorescence could only be correctly assigned after subtracting the blue autofluorescence background. The plaque cores have very strong blue autofluorescence which is roughly 2.09 times of average DAPI nuclear staining and roughly 1.78 times of average Hoechst nuclear staining. The composite flower-like structures formed by Cathepsin D lysosomal staining wrapping dense core blue fluorescence should not be considered as central-nucleated neurons filled with defective lysosomes since there was no nuclear staining in the plaque core when the blue autofluorescence was subtracted. Furthermore, the dense cores were shown to be completely lack of nuclear signals by PI staining. The Aβ aggregation assay indicated that both Aβ self-oligomers and Aβ/Hemoglobin (Hb) heterocomplexes had significant blue autofluorescence. However, the blue autofluorescence intensity was not always proportional to the intensity of Aβ immunostaining. The majority of aggregates in the Aβ/Hb incubations were sensitive to Proteinase K (PK) digestion while the rest were PK resistant. The blue autofluorescence of Aβ aggregates not only labels senile plaques but also illustrates red blood cell aggregation, hemolysis, CAA, vascular amyloid plaques, vascular adhesion and microaneurysm. In summary, we conclude that Aβ-aggregation-generated blue autofluorescence is an excellent amyloid pathology marker in the senile plaques, blood and vascular pathologies in the Alzheimer’s disease.

## Introduction

Senile plaques are the most important hallmark of Alzheimer’s disease, with amyloid beta (Aβ) as a major biochemical component in senile plaques. Senile plaque blue autofluorescence, which we assigned a temporary name of “MetaBlue”, is another interesting character of senile plaques, which has been investigated by several studies including our own[1-4]. An early study identified the blue autofluorescence in senile plaques in AD around 40 years ago[1]. More recent studies showed that, besides senile plaques, blood vessels with cerebral amyloid angiopathy (CAA) similarly had this blue autofluorescence[2, 3]. Our studies further showed that some populations of red blood cells (RBCs) also had strong blue autofluorescence[4]. In addition, we showed that RBC blue autofluorescence is associated with increased calcium level and Aβ content. Although senile plaque blue autofluorescence is now a well-known phenomenon, its impact on AD pathogenesis and AD research was not fully addressed.

We analyzed senile plaque blue autofluorescence with immunohistochemistry and fluorescence imaging on AD patient frontal brain sections and also analyzed the Aβ aggregation process *in vitro* in test tubes in order to find out whether the blue autofluorescence is coupled to the Aβ aggregation process. The results showed that the Aβ peptide aggregation directly contributes to the senile plaque blue autofluorescence and the amyloid blue autofluorescence could be a valuable label-free marker for both senile plaques and a spectrum of important blood and vascular pathologies in the Alzheimer’s disease.

## MATERIAL AND METHODS

### Tissue sections

Alzheimer’s disease patient frontal lobe brain paraffin tissue sections were purchased from GeneTex (Irvine, CA, USA). Additionally, AD patient frontal lobe brain paraffin tissue sections were provided by National Human Brain Bank for Development and Function, Chinese Academy of Medical Sciences and Peking Union Medical College, Beijing, China. This study was supported by the Institute of Basic Medical Sciences, Chinese Academy of Medical Sciences, Neuroscience Center, and the China Human Brain Banking Consortium. All procedures involving human subjects are done in accord with the ethical standards of the Committee on Human Experimentation in Shanghai Jiao Tong University and in Xinhua Hospital, and in accord with the Helsinki Declaration of 1975.

### List of antibodies for immunohistochemistry (IHC)

The following primary antibodies and dilutions have been used in this study: Aβ/AβPP (CST #2450, 1:200), Cathepsin D (Abcam ab75852, 1:200), HBA (Hemoglobin alpha chain) (Abcam ab92492, 1:200). The following secondary antibodies and dilutions have been used in this study: donkey anti-mouse Alexa-594 secondary antibody (Jackson ImmunoResearch 715-585-150, 1:400), donkey anti-rabbit Alexa-488 secondary antibody (Jackson ImmunoResearch 711-545-152, 1:400), donkey anti-rabbit Alexa-594 secondary antibody (Jackson ImmunoResearch 711-585-152, 1:400) and donkey anti-mouse Alexa-488 secondary antibody (Jackson ImmunoResearch 715-545-150, 1:400).

**Immunohistochemistry** was performed as described[5]. Briefly, paraffin sections were firstly deparaffinized by Xylene, 100% EtOH, 95% EtOH, 75% EtOH, 50% EtOH, and PBS washes. Sections were then treated with 10 mM pH6.0 sodium citrate or 10mM pH9.0 Tris-EDTA antigen retrieval solutions with microwave at high power for 5 minutes to get to the boiling point and then at low power for another 15 minutes. The sections were allowed to naturally cool down to room temperature. Then, the slides were blocked with TBST+3% BSA solution for 1 hour at room temperature. After blocking, the samples were incubated with primary antibodies at room temperature for 2 hrs followed by 5 washes of TBST. After that, the samples were incubated with fluorescent secondary antibodies overnight at 4 °C. The treated samples were washed again with TBST 5 times the second day and mounted with PBS+50% glycerol supplemented with DAPI (Sigma, D9542, 1 μg·mL^−1^, St. Louis, MO, USA) or H33342 nuclear dye (Sigma; B2261, 1 μg·mL^−1^) and ready for imaging. Both DAPI and H33342 stained the nuclei with blue fluorescence. To differentiate nuclear staining from senile plaque blue autofluorescence signal, in some of the IHC experiments, PI (Sigma; P4170, 10 μg·mL^−1^) was used as a red nucleus fluorescence dye instead. To analyze the blue autofluorescence in those IHC experiments with also green and red fluorescent secondary antibodies, no nuclear dye was included. IHC experiments without primary antibodies were used as negative controls. All experiments have been repeated twice taking from two different AD brain samples in order to verify the reproducibility of the results.

**Slide-based Aβ aggregation assay** was performed as described[4] with slight modifications. Briefly, human Aβ40 or Aβ42 peptides (Apeptide Co., Shanghai, China) were first solubilized with DMSO at 1 mg/mL concentration. Human Hemoglobin (Hb) (H7379, Sigma) was solubilized in sterile H_2_O at 10 mg/mL concentration. Then, Aβ peptides were diluted into sterile PBS solution at 5 μM concentration with or without 1.25 μM Hemoglobin. The mixture was incubated at 37 degree up to 3 days. Samples were obtained at Day 0, Day 1, Day 2, and Day 3. 1 μl out of each obtained sample was spotted onto adhesive glass slides and dried up in a 37-degree incubator. The glass slides with samples were fixed with 4% formaldehyde in PBS for 10 minutes at room temperature and washed 3 times with PBS. To test the sensitivity of aggregates to protease digestion, paralleled slides were treated with Proteinase K at 20 μg/mL in 10 mM TE (pH 7.5) buffer for 15 minutes at room temperature. The procedure to analyze the Aβ aggregates on the slides using antibodies was the same as in immunohistochemistry. Since our previous experiments showed very similar Aβ complex formation in samples obtained from Day 0 to Day 3[4], only Day 3 samples were shown in this study. All experiments have been repeated in order to verify the reproducibility of the results.

### Imaging analysis

The fluorescent images were taken with a CQ1 confocal fluorescent microscope (Yokogawa, Ishikawa, Japan). The blue autofluorescence of senile plaques (MetaBlue) can be imaged with the standard DAPI blue fluorescence channel using the CQ1 confocal microscope with the excitation wavelength of 405 nm,and the emission spectrum of 447 nm-460 nm. When comparing the blue fluorescence of senile plaques without or with nuclear dye staining, the same senile plaques were imaged twice with identical imaging conditions, once without dye staining with only blue autofluorescence and the second time with nuclear dye staining by DAPI or Hoechst. Images were then analyzed with Image J software (a free imaging analysis software from NIH, USA). Mean area density was used as the parameter to define the marker densities. The perimeters of different structures were measured to simplify the diameter estimation by using the following formula: The diameter= the perimeter/3.14. The autofluorescence signal subtraction was performed with the Image J software with the image calculation toolbar. When performing the autofluorescence subtraction analysis, the subtraction was only carried out for the blue channel images.

### Statistics

All data first went through a Shapiro & Wilk normality test using SPSS Statistics 19 software. Two-tailed unpaired T-test was used to compare the means of data with a normal distribution with Excel 2007 software. For data that were not normally distributed, nonparametric Kruskal-Wallis test was performed to compare the means of unpaired samples and Wilcoxon Signed Rank test was applied to compare the means of paired samples by using SPSS Statistics 19 software. The p Value threshold for statistical significance is set at 0.05. If p<0.05, then the difference was considered statistically significant. When p<0.001, it was labeled as p<0.001. If p≧0.001, then the p Value was shown as it was.

## Author contributions

HF conceived the study, designed and supervised the experiments and wrote the manuscript; HF and JL performed the experiments; HF, JL, CZ, PD, GG, QG, XG and DC did the data analysis; all authors reviewed the manuscript.

## Data availability statement

The data that supports the findings of this study are available from the corresponding author upon request.

## Acknowledgements

We want to thank the help from Dr. Ma Chao and Dr. Qiu Wenying for providing AD tissue sections from National Human Brain Bank for Development and Function, Chinese Academy of Medical Sciences and Peking Union Medical College, Beijing, China. Additionally, we want to thank Housheng Wang, Pingxin Liu, Xiaoyi Bao, and Jun Wang for excellent lab assistance.

## Results

### The blue fluorescence of the cores of dense-core plaques is stronger than nuclear staining by blue fluorescent nuclear dyes such as DAPI and Hoechst

Although it is well-known that senile plaques have intrinsic blue fluorescence, it is not clear how strong it is compared to conventional nuclear staining. Previously, we analzyed senile plaque blue fluorescence with or without Hoechst nuclear staining[4]. In this study, we also analyzed senile plaque blue fluorescence with or without DAPI nuclear staining (Figure 1A, Figure 1B). We focused the fluorescence intensity analysis on the plaque cores of dense-core senile plaques since they had the strongest blue autofluorescence. The average plaque core blue fluorescence intensity with DAPI staining is 1.03±0.03 times of the average plaque core blue fluorescence density without DAPI staining (12 plaques measured, p=0.024 by Wilcoxon Signed Rank Test). Although there might be a little increase of the fluorescence intensity by DAPI staining of the plaque cores but the difference is not large, the bulk of blue fluorescence in the plaque cores is likely due to the blue autofluoresence. When we substracted the senile plaque blue autofluorescence from the DAPI or Hoechst-stained image, the correct identification of DAPI or Hoechst-stained nuclei could be achieved and there was no nuclear staining in the plaque cores (Figure 1A bottom panel and Figure 1B bottom panel).

**Figure 1:**
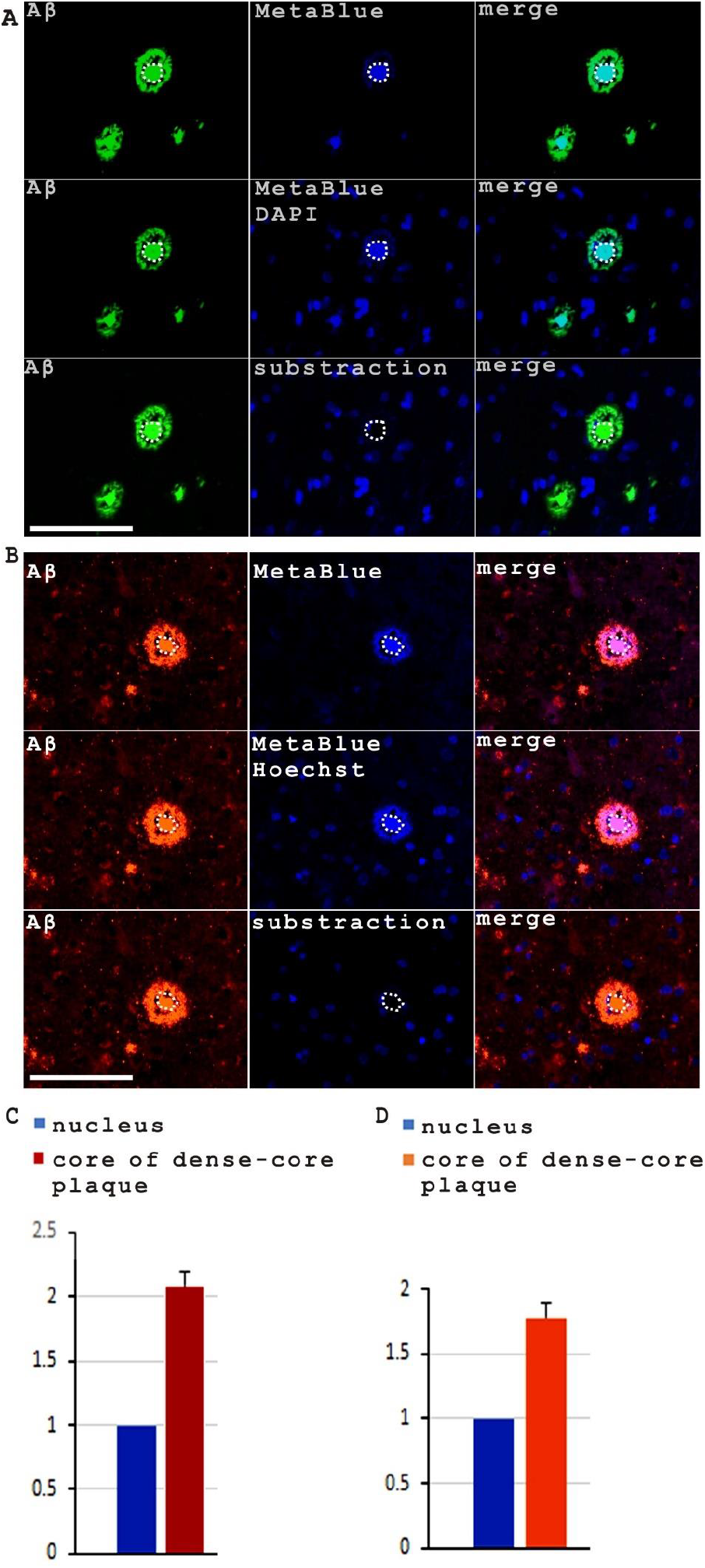
The blue autofluorescence of the cores of dense-core senile plaques is stronger than conventional blue fluorescent nuclear staining. (**A**) An example image showed the plaque core blue fluorescence of a dense-core plaque without (top panel) or with DAPI staining (middle panel). When the senile plaque blue autofluorescence was subtracted from the DAPI stained image, then the correct identification of DAPI-stained nuclei was shown (bottom panel). The dashed circle indicated the plaque core. Scale bar, 100 μm. (**B**) An example image showed the plaque core blue fluorescence of a dense-core plaque without (top panel) or with Hoechst (H33342) nuclear staining (middle panel). When the senile plaque blue autofluorescence was subtracted from the Hoechst-stained image, then the correct identification of Hoechst-stained nuclei was shown (bottom panel). The dashed circle indicated the plaque core. Scale bar, 100 μm. (**C**) The intensity of plaque core blue autofluorescence is around 2.09 (±0.10) times of DAPI nuclear staining of brain tissue cells. (**D**) The intensity of plaque core blue autofluorescence is around 1.78 (±0.11) times of Hoechst nuclear staining of brain tissue cells.

We then measured the plaque core blue fluoresence intensity comparing to the average nuclear fluoresence intensity. The results showed that the cores of dense-core plaques (N=11) have much stronger blue fluorescence which is roughly 2.09 (±0.1) times of average DAPI nuclear staining (11 plaque cores vs 110 cell nuclei) and roughly 1.78 (±0.11) times of average Hoechst nuclear staining (10 plaque cores vs 100 cell nuclei) (Figure 1C, 1D), indicating that the blue fluorescence of the cores of dense-core plaques is significantly stronger than average nuclear staining by DAPI (P<0.001, Kruskal-Wallis test) and Hoechst staining (P<0.001, Kruskal-Wallis test). We also measured the average diameter of the cores of dense-core plaques (15.84±2.80 μm, N=11) while the average diameter of the nuclei of brain tissue cells is 9.70±1.49 μm (N=110). The diameter of the cores of dense-core plaques is 1.63±0.29 times of the average diameter of brain tissue cell nuclei (p<0.001, T-test).

### The cores of dense-core plaque also co-localized with Cathepsin D staining along with blue autofluorescence

Our previous research showed that many senile plaques come with a diffusive staining of Cathepsin D, which is quite different from the strong granule-like lysosomal Cathepsin D staining from surrounding neural cells[4]. In this study, we checked the co-expression of senile plaque Aβ, cathepsin D and blue auto fluorescence. The experiment showed that the cores of dense-core plaques were stained for all three markers (Figure 2). Flower-like composite structures were formed by abundant lysosome vesicles surrounding the central plaque cores with the strong blue fluorescence. Although these flower-like structures resembled neural cells in appearance (Figure 2B) but they were certainly not central-nucleated neurons that were filled with defective lysosomes because the bulk of blue fluorescence in the plaque cores derived from senile plaque blue autofluorescence but not from nuclear staining. When we substracted the blue autofluorescence from the Hoechst-stained image, then correct identification of Hoechst-stained nuclei could be achieved, which showed that the plaque core was lack of nuclear staining by Hoechst while true nucleated cells retained the blue nuclear fluorescence (Figure 2C).

**Figure 2.**
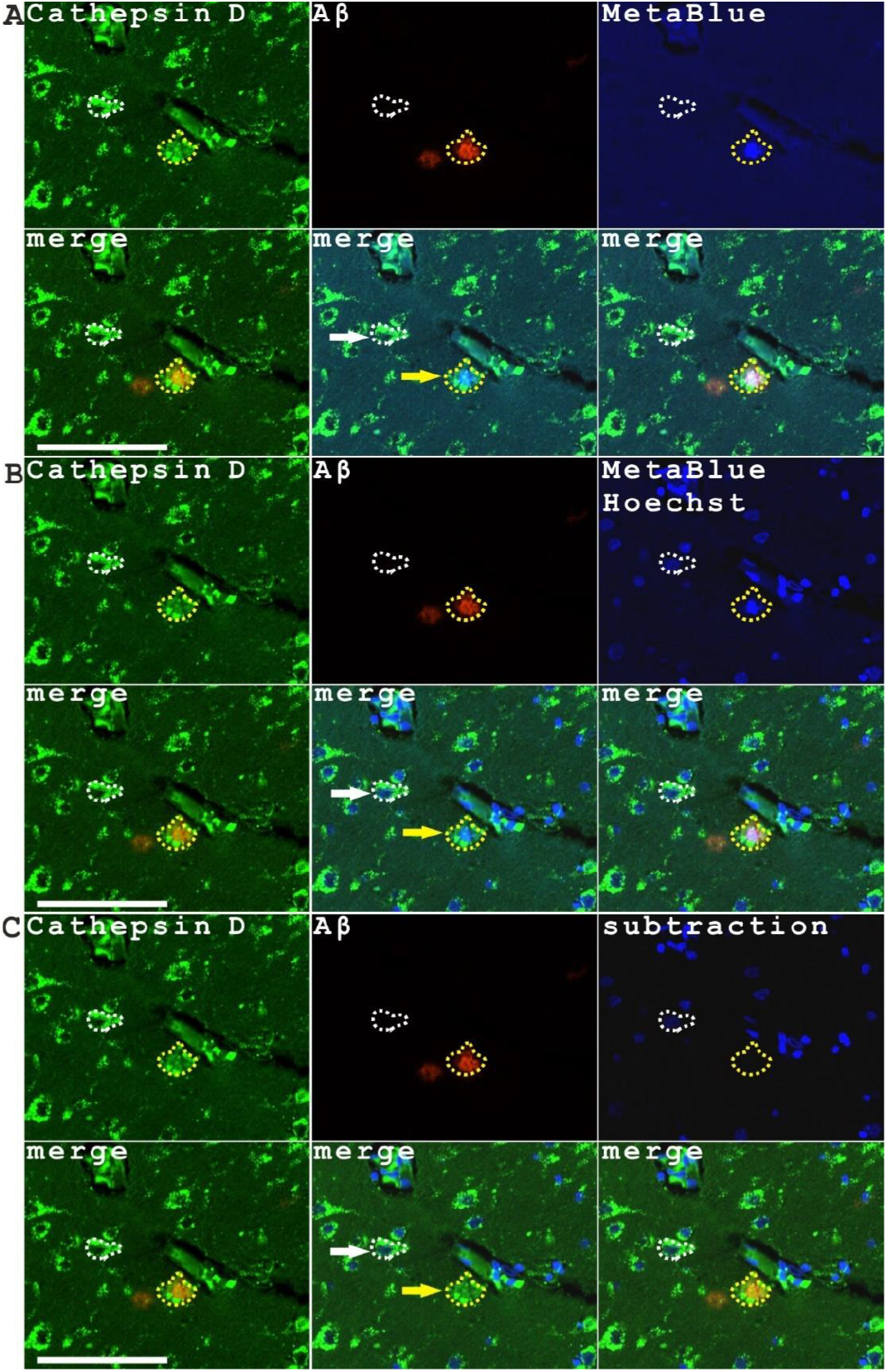
The cores of dense-core plaque also colocalized with Cathepsin D staining along with blue autofluorescence. An example image showed senile plaque Cathepsin D, Aβ and blue fluorescence without (**A**) or with Hoechst nuclear staining (**B**). When the senile plaque blue autofluorescence was subtracted from the Hoechst-stained image, then the correct identification of Hoechst-stained nuclei was shown (**C**) while the senile plaque blue autofluorescence was completely removed. The dashed yellow lines circled the dense-core plaque along with dense peri-plaque lysosomal Cathepsin D signals, indicating the composite structure with a plaque core wrapped with Cathepsin D staining. The dashed white lines indicated a nucleated neural cell. The yellow and white arrows highlighted the merged images from the green and blue channels for the plaques and nucleated neural cells. Although these two structures look alike by appearance, but they are definitely different structures since the blue fluorescence in neural cells coming from the nuclear staining while the blue fluorescence of senile plaques comes from senile plaque autofluorescence. Scale bars, 100 μm.

### The cores of dense-core plaques were completely lack of nuclear staining by PI

Since nuclear staining is an important aspect of pathological studies in Alzheimer’s disease, it is important to identify nuclei correctly and to differentiate senile plaque autofluorescence from nuclear signals. We noticed that senile plaques have strong blue autofluorescence but weak signals in the red channel so that we switched to using a red fluorescent nuclear dye, PI, to identify cell nuclei on AD pathological sections. Our results showed all dense-core plaques have strong blue autofluorescence and Aβ staining but are completely lack of nuclear staining by PI (Figure 3A, B, C). The figure also showed a senile plaque with a central nucleus labeled by PI but this plaque was lack of blue autofluorescence and Aβ staining in the plaque core thus should not be regarded as a dense-core plaque (Figure 3D).

**Figure 3.**
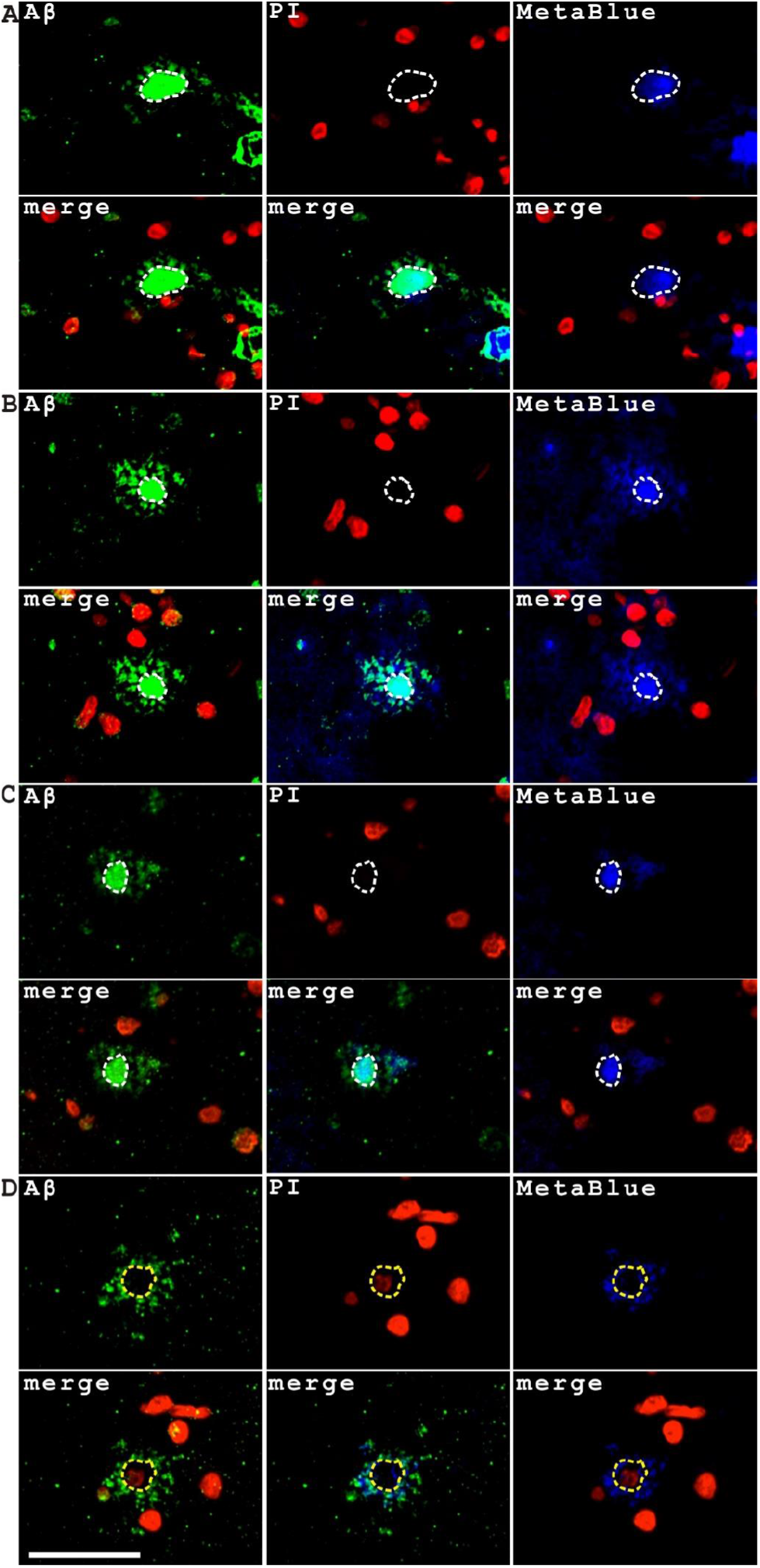
The cores of dense-core plaques with strong blue autofluorescence and Aβ staining do not have nuclear staining by the red fluorescence nuclear dye PI. **(A, B, C**) showed three dense-core senile plaques with their cores labeled by both Aβ antibody and MetaBlue but lack of nuclear staining. The dashed circle indicated the plaque cores. (**D**) showed a plaque with peripheral Aβ staining and a central portion lack of Aβ staining but with nuclear staining by PI, however, this plaque is not a dense-core senile plaque. The yellow dashed circle indicated the central region of the plaque. Scale bar, 50 μm.

### Aβ aggregation process directly contributes to the blue autofluorescence generation

The mechanisms producing senile plaque blue autofluorescence are not very clear. Previously we developed an *in vitro* assay to examine the amyloid peptide aggregation process on adhesive slides[4], we re-did the aggregation experiments and examined the changes of amyloid blue fluorescence during the aggregation process. The experiment showed that the Aβ aggregation indeed came together with a characteristic blue autofluorescence (Figure 4). Aβ40 self-oligomers showed significant blue autofluorescence while the majority of Aβ42 self-oligomers showed weak blue autofluorescence signals (Figure 4A) with only a few Aβ42 aggregate patches showing similar intensity blue autofluorescence as Aβ40 self-oligomers (Figure 4B). However, both Aβ40/Hb hetero-oligomers and Aβ42/Hb hetero-oligomers have clear blue autofluorescence signals (Figure4C, 4D). We also noticed that the intensity of blue autofluorescence is not always proportional to the intensity of Aβ immunostaining, which suggests that the yield of blue autofluorescence is probably more affected by the configuration and structure property of oligomers rather than the quantity of amyloid peptides. This experiment suggested that Aβ aggregation directly contributes to “MetaBlue” autofluorescence.

**Figure 4.**
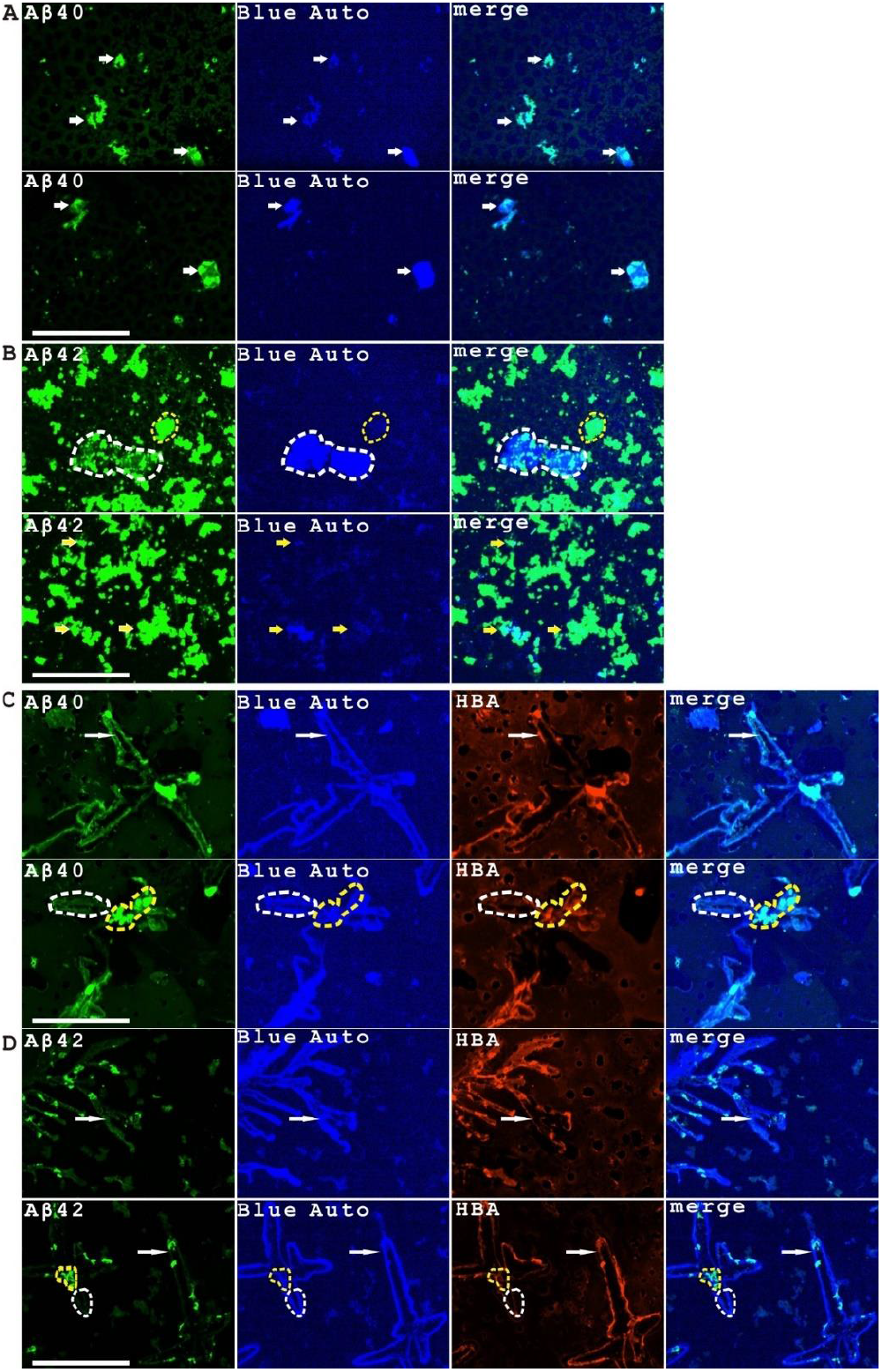
Aβ aggregation process is intrinsically associated with blue autofluorescence production. (**A**) Aβ40 self-oligomers showed significant blue autofluorescence (labeled in short as “Blue Auto”) as indicated by arrows. (**B**) Aβ42 self-oligomers showed two kinds of patterns of blue autofluorescence with a few strong patches while majority of the aggregates have very weak blue autofluorescence. A strong intensity patch was indicated by the white dashed lines. Other structures with weak blue fluorescent signals were indicated by yellow dashed lines or arrows. (**C**) Aβ40/Hb heterocomplex aggregates showed significant blue autofluorescence as indicated by arrows. However, the blue fluorescence intensity was not correlated to Aβ immunostaining intensity, showing by the comparison between the structures indicated by white dashed or yellow dashed lines. These regions had very different intensity of Aβ immunostaining but similar intensity of blue autofluorescence. (**D**) Aβ42/Hb heterocomplex aggregates also showed significant blue autofluorescence as indicated by arrows. Again, the blue fluorescence intensity was not correlated to Aβ immunostaining intensity, showing by the comparison between the structures indicated by white dashed or yellow dashed lines. These regions had very different intensity of Aβ immunostaining but similar intensity of blue autofluorescence. Scale bars, 100 μm.

Aβ aggregates are often considered as toxic aggregates that resists proteinase digestion. However, it is possible some Aβ aggregates have physiological functions and could be metabolically degraded. We checked the proteinase sensitivity of Aβ aggregates with blue auto fluorescence by Proteinase K digestion. The data showed that the hetero-oligomers formed by Aβ40/Hb or Aβ42/Hb complexes were Proteinase K sensitive, thus could possibly be physiologically degraded (Figure 5A top panel, Figure 5B top panel), while some Aβ40 or Aβ42-dominant complexes were resistant to PK digestion and still preserving blue auto fluorescence after proteinase treatment (Figure 5A bottom panel, Figure 5B bottom panel).

**Figure 5.**
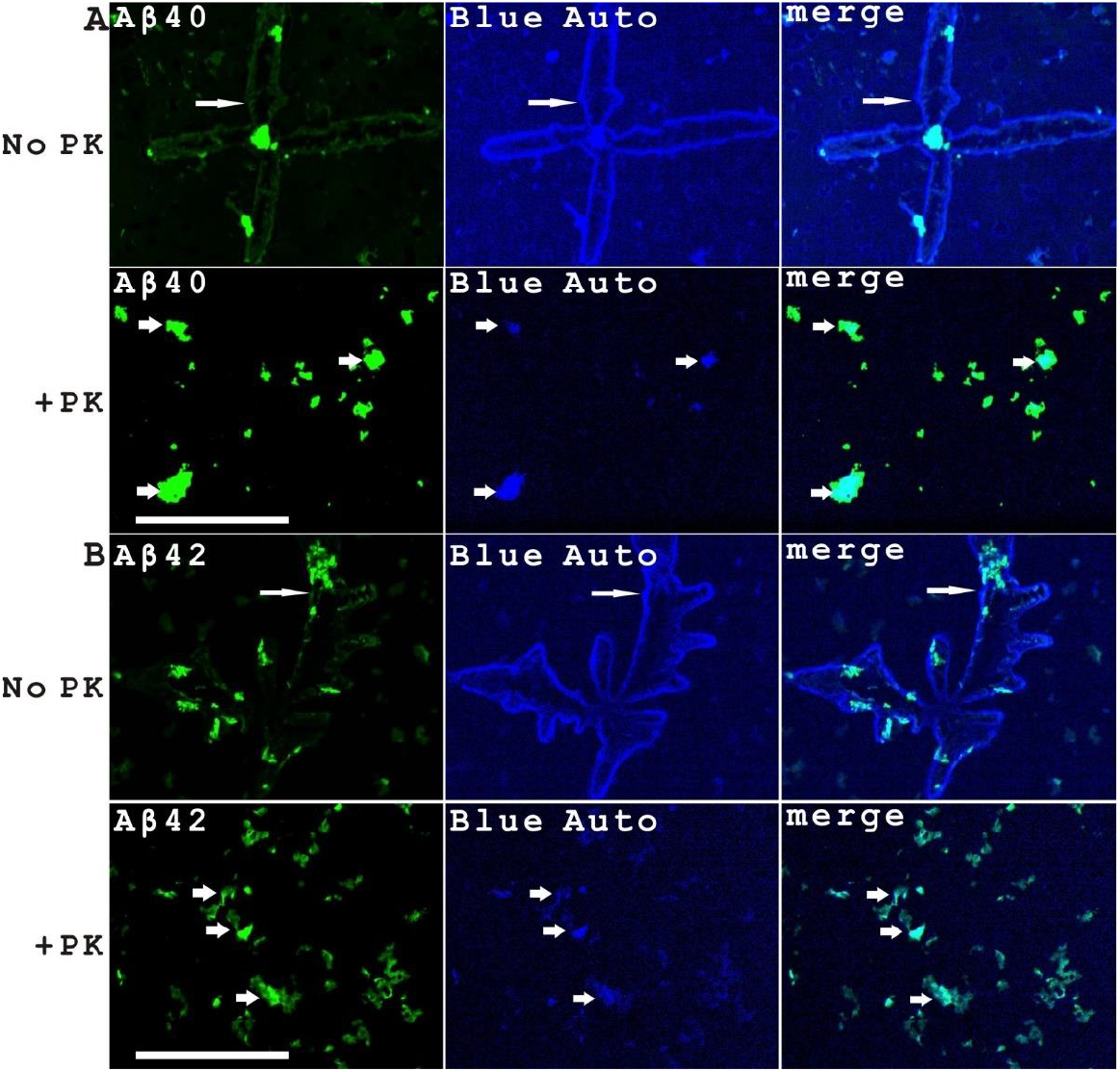
The majority of blue fluorescent Aβ aggregates formed during Aβ/Hb incubation were sensitive to Proteinase K digestion while the rest were resistant to Proteinase K digestion. (**A**) The image showed the large structures formed in the Aβ40/Hb incubation were sensitive to PK digestion while some smaller amyloid aggregates were resistant to PK digestion. (**B**) The image showed the large structures formed in the Aβ42/Hb incubation were sensitive to PK digestion while some smaller amyloid aggregates were resistant to PK digestion. Scale bars, 100 μm.

### MetaBlue is an excellent label-free marker of multiple blood and vascular pathology hallmarks besides senile plaques in the Alzheimer’s disease

Although senile plaques and neurofibrillary tangles are considered as the main pathological hallmarks in Alzheimer’s disease, many studies indicated that multiple blood and microvessel defects are also important pathological phenotypes in AD[4, 6-10]. We studied the Aβ staining and blue autofluorescence in multiple blood and vascular pathologies in AD brain tissues as shown in Figure 6. The data showed that MetaBlue is associated with Aβ expression in aggregating RBCs and in hemolysis (Figure 6A, B). MetaBlue also associated with Aβ expression in all CAA blood vessels we examined (Figure 6C, D, E). Aggregating RBCs (Figure 6C) and transparent lumen-blocking vascular amyloid plaques with MetaBlue fluorescence (Figure 6D) were sometimes observed in the CAA blood vessels. Many CAA vessels had no blood flow (Figure 6E showed one such example), suggesting both hemolysis and blocking of blood flow could be happening. Additionally, in some CAA vessels, vascular adhesion and lumen closure phenotype was observed (Figure 6F). MetaBlue is also a high-contrast marker of microaneurysms in AD tissues (Figure 6G, H). All these results indicated that MetaBlue is a good marker of blood and vascular abnormalities comparable to Aβ immunostaining with high contrast and sensitivity.

**Figure 6.**
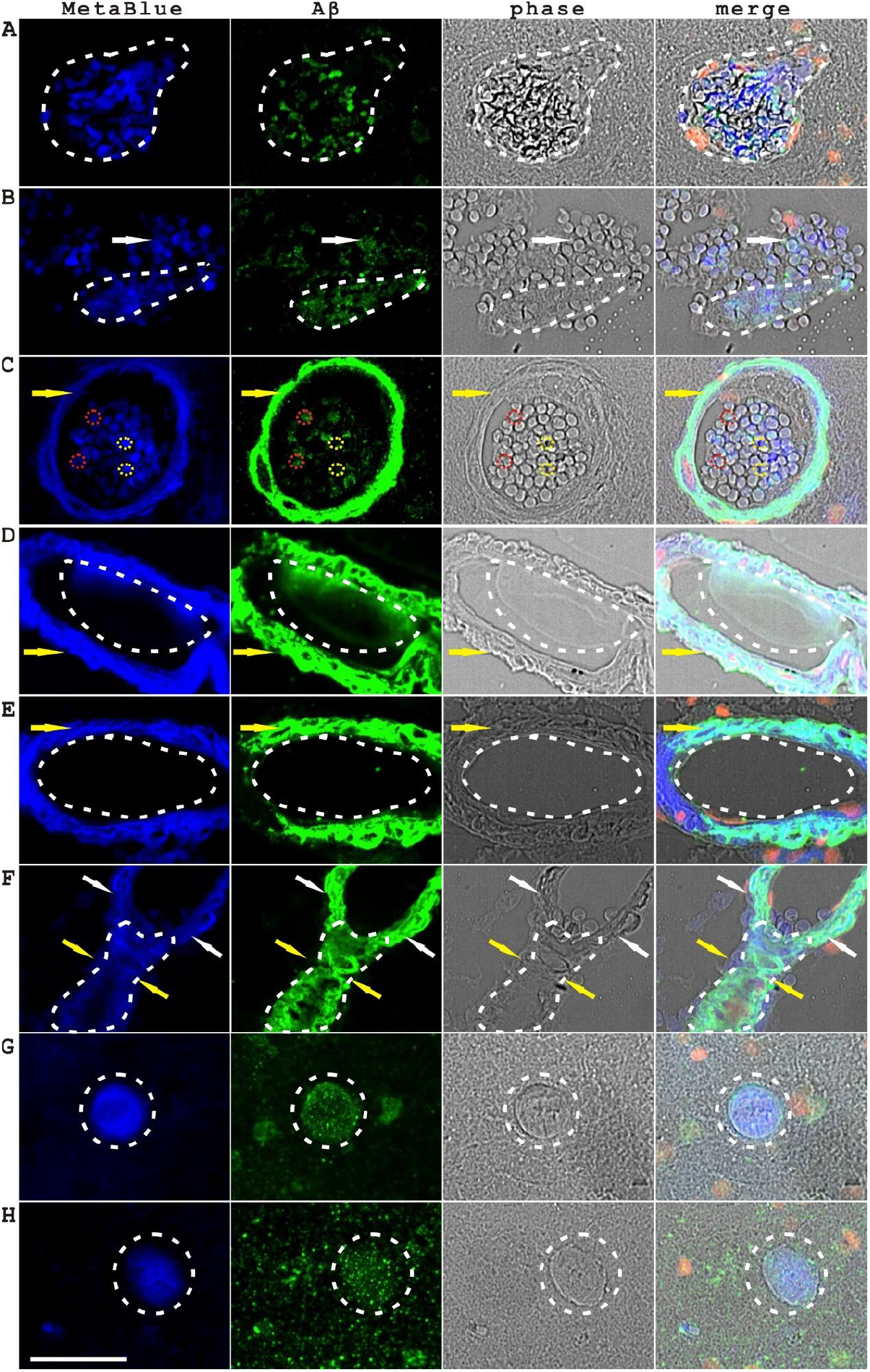
As an intrinsic indicator of Aβ aggregation, MetaBlue (Aβ amyloid blue fluorescence) is a good label-free marker of multiple complicated blood and vascular pathologies in Alzheimer’s disease. (**A**) MetaBlue associates with Aβ expression in abnormally aggregating red blood cells within a blood vessel also showing vascular wall protrusions, indicated by dashed lines. (**B**) MetaBlue associates with Aβ expression in aggregating red blood cells (indicated by arrows) and also in hemolytic RBCs (indicated by dashed lines). (**C**) MetaBlue associates with Aβ expression in CAA (indicated by arrows) and in aggregating RBCs (indicated by dashed circles). At single cell levels, the blue autofluorescence intensity of red blood cells was not necessarily correlated to the Aβ staining intensity. The red dashed circles showed two RBCs with relatively high Aβ intensity but weaker blue fluorescence comparing to two RBCs indicated by yellow circles with relatively low Aβ intensity but stronger blue fluorescence. (**D**) MetaBlue associates with Aβ expression in CAA (indicated by arrows) and in vascular amyloid plaques (indicated by dashed lines). The indicated vascular amyloid plaque is blocking the major part of the vascular lumen. (**E**) MetaBlue associates with Aβ expression in a CAA blood vessel (indicated by arrows) without blood flow (indicated by dashed lines, showing the empty vessel lumen without any red blood cells). (**F**) MetaBlue associates with Aβ expression in a CAA blood vessel showing a vascular adhesion phenotype (indicated by dashed lines) with an abrupt closure (indicated by yellow arrows) of the blood vessel lumen (the open blood vessel lumen was indicated by white arrows). (**G-H**) MetaBlue associates with Aβ expression in two examples of microaneurysm (indicated by dashed circles) in AD tissues. Scale bar, 50 μm.

## Discussion

Senile plaque blue autofluorescence is a well-known character of senile plaques. However, senile plaque blue autofluorescence signals were often ignored in AD studies. We did the side-by-side comparisons of senile plaques stained with or without blue nuclear dyes such as DAPI and Hoechst with standard nuclear staining conditions. The cores of dense core plaque have very strong blue fluorescence which is roughly 2.09 times of average DAPI nuclear staining and roughly 1.78 times of average Hoechst nuclear staining. This result indicated that plaque core blue fluorescence yielded stronger signals than conventional blue nuclear dye staining that should not be ignored in pathological studies.

We also analyzed senile plaque blue autofluorescence in the context of Cathepsin D and Aβ co-immunostaining. The cores of dense core plaques were enriched with all three markers, indicating that Cathepsin D might be related to senile plaque formation. The flower-like pattern of lysosomal Cathepsin D staining wrapping central plaque blue autofluorescence is almost identical with or without Hoechst nuclear dye staining, indicating these structures are senile plaques surrounded by lysosomal markers and they are not central-nucleated neurons that are filled with defective lysosomal vesicles. The correct identification of cellular nuclei could be achieved by subtracting the blue autofluorescence from DAPI or Hoechst-stained images, which also showed that the senile plaque blue fluorescence mostly came from blue autofluorescence but not blue nuclear dye staining. Additionally, the nuclei-null property of dense cores could be proved by the red fluorescence nuclear dye PI staining. PI staining results verified that all the cores of dense-core plaques do not have PI-labeled nuclei.

Several mechanisms might cause senile plaque blue autofluorescence. It could be due to the intrinsic property of aromatic amino acids such as tryptophan, tyrosine and phenylalanine[11], or because of aromatic amino acid oxidation modifications[12] or could be due to the presence of other blue fluorescent materials in the senile plaques[13, 14]. A simpler explanation could be that the blue autofluorescence is induced by the amyloid peptide aggregation process as suggested from a previous study[15]. The *in vitro* amyloid aggregation experiment showed that the Aβ aggregation process indeed produced blue autofluorescence. Aβ40 self-aggregates are often blue fluorescent while Aβ42 self-aggregates were less-frequently blue fluorescent. Aβ40/Hb incubation as well as Aβ42/Hb incubation produced heterocomplexes structures with significant blue fluorescence, which were sensitive to PK digestion. The blue autofluorescence of heterocomplex of Aβ and Hemoglobin also explained the blue fluorescence present in RBCs showing stressed phenotypes since Aβ might interact with Hb in RBCs under stress conditions. In our experiments, the intensity of blue autofluorescence was not always proportional to the intensity of Aβ immunostaining as shown in Aβ42 oligomers, Aβ40/Hb heterocomplex aggregates, Aβ42/Hb heterocomplex aggregates and even in individual red blood cells, which might suggest that the yield of blue fluorescence is a property of amyloid aggregate structure or configuration instead of the absolute amount of Aβ peptides. The specific Aβ oligomer subtypes that are responsible for the high-efficiency blue autofluorescence generation need to be studied further. Our previous experiments showed that Aβ and Hemoglobin co-localize in both in vitro experiments and on pathological sections, suggesting that Aβ aggregation process was happening early in stressed red blood cells during AD development[4]. Most of these amyloid aggregates might be metabolically degraded given their sensitivity to PK digestion.

It is important to recognize that the pathological hallmarks in Alzheimer’s disease are not limited to the senile plaques and neurofibrillary tangles, but also included multiple important blood and vascular pathologies, such as red blood cell aggregation, intravascular hemolysis, CAA, vascular amyloid plaques, vascular adhesion and microaneurysms, associating with both Aβ aggregation and aggregation-associated blue autofluorescence as emphasized in the current study. Cerebral hypoperfusion induced by vascular defects has long been hypothesized as the culprit of Alzheimer’s disease[16, 17]. Several studies linked the brain capillary abnormalities closely to AD pathogenesis[7, 10, 18, 19]. Recent publications also demonstrated that blood-borne Aβ contributes to Alzheimer’s disease pathologies[20, 21]. Microaneurysm rupture and the chronic leakage of Aβ aggregates might be the potential mechanism of senile plaque formation[4]. Whether the Aβ amyloid blue fluorescence could be a sensitive, label-free, low cost and convenient marker for the *in vivo* monitoring of the amyloid pathology of senile plaques, blood and vascular defects of AD patients is an interesting subject for the future investigation.

